# BMP signaling underlies the craniofacial heterochrony in phyllostomid bats, a hyperdiverse mammal group

**DOI:** 10.1101/2021.05.17.444516

**Authors:** Jasmin Camacho, Jacky D. Lin, Michaela McCormack, Rachel Moon, Samantha K. Smith, John J. Rasweiler, Richard R. Behringer, Clifford J. Tabin, Arhat Abzhanov

## Abstract

The potential for variation and the capacity to evolve in response to ecological opportunity are important aspects of an adaptive radiation. Identifying the origin of phenotypic variation, in which natural selection might act upon, is a major goal of evolutionary developmental biology. The New World leaf-nosed bats (phyllostomids) are a textbook example of an adaptive radiation. Their cranial morphology is diverse along relative facial length, which is related to their diets. We previously used geometric morphometrics to reveal peramorphosis, a type of heterochrony, in the cranial evolution among phyllostomid bats. We then demonstrated that the mechanism of peramorphic diversity in phyllostomid rostrum length resulted from altered cellular proliferation. Here, we investigate the progenitors of the face, the cranial neural crest, and a key signaling pathway related to their proliferation and differentiation into mature tissues: the bone morphogenetic protein (BMP). With geometric morphometrics, immunofluorescence, and confocal imaging—in three phyllostomid species and one outgroup bat species—we show the molecular patterns that underlie the adaptive and innovative traits seen in phyllostomid bats. Then, with mouse genetics, we mimic the BMP molecular pattern observed in nectar feeding bats and recapitulate the elongated morphological variation in mice. Surprisingly, we also observe an expansion in the nose-tip of mice, akin to the expanding leaf-nose tissue in phyllostomid bats. These data, combined with the mouse genetics literature on BMP signaling, suggest the BMP developmental pathway plays a central role in shaping the craniofacial variation necessary for adaptation in bats. Further, we speculate that the BMP signaling pathway could underlie other bizarre facial phenotypes in mammals that are derived from frontonasal mesenchyme, such as the proboscis. Overall, this study combines a comparative framework to developmental data, with a genetic approach, to directly investigate the role of development on complex morphology.

## Introduction

### Mammal craniofacial skeleton

The mammalian craniofacial skeleton is one of the most morphologically diverse structures in the vertebrate body and among mammalian groups. It has adaptations for hearing, protection of the brain, defensive armor, feeding performance, and many other tasks. Best representing this diversity are the New World leaf-nosed bats (Order Chiroptera, Family Phyllostomidae). They have a broad range of morphologies associated with dietary specialization (Freeman 2000; Van Cakenberghe et al. 2002; Baker et al. 2012; Rossoni et al. 2017). Primarily, their cranial morphology is specialized to diet along the relative facial length of the head (Freeman 2000; Dumont et al. 2014; Sears 2014; Sorensen et al. 2014; Arbour et al. 2019; Hedrick et al. 2019; Rossoni et al. 2019). The facial length differences between species evolved from a heterochronic shift called peramorphosis to extend the period of craniofacial growth (Camacho et al. 2019) and the peramorphic changes in phyllostomid facial growth arise from altered cellular proliferation (Camacho et al. 2020). The genetic mechanism underlying this heterochronic alteration is unknown; however, studies using mouse genetics reveal hypothetical cellular mechanisms generating facial variation.

### Cellular basis of facial length differences

There are several plausible developmental cell mechanisms of cranial neural crest (CNC) and their derivatives that influence variation in facial length development including migration, proliferation, and survival. Since we recently demonstrated that the mechanism of peramorphic diversity in phyllostomid rostrum length resulted from altered cellular proliferation (Camacho et al. 2020), we will focus on this cellular mechanism. From mouse genetics, proliferation of the ectomesenchyme (CNC-derived mesenchymal tissue) has been shown to affect facial length through establishing condensations, which are related to the number of committed precursor cells that will give rise to cartilage and/or bone (Helms and Schneider 2003; Akiyama et al. 2005; Chai and Maxson 2006). Another widespread occurrence affecting facial length is proliferation within the cartilage or bone lineage (Pavlov et al. 2003; Akiyama et al. 2005; Brugmann et al. 2007; Nagayama et al. 2008; Kaucka et al. 2016). These cellular types may be distinguished using known molecular markers, such as ectomesenchyme cells expressing PAX7 (Murdoch et al. 2012), pre-chondrogenic (cartilage) progenitor cells expressing SOX9 (Bell et al. 1997), pre-osteoblast (bone) progenitor expressing RUNX2 (Chen et al. 2012), mature chondrocyte cells expressing SOX9 and COL2 (Mori-Akiyama et al. 2003), and mature osteoblast cells expressing COL1 and OSX (Komori 2010).

### CNC regulation

To date, the facial development underlying evolutionary craniofacial variation in phyllostomids has been based on the morphological analysis of midfacial length after the fusion of the craniofacial prominences (~Carnegie Stage 16) and until sutural fusion as pups. These studies have shown that distinct species of phyllostomids range in craniofacial length beginning at Carnegie Stage 18 (Sears 2014; Camacho et al. 2019), with evolutionary changes to facial length correlating with cellular proliferation changes to the midface (Camacho et al. 2020). Carnegie Stage (CS)18, which is approximately equivalent to mouse E12.5 based on limbs (Cretekos et al. 2005), is a critical period in midfacial morphogenesis associated with skeletal tissue development (Ishii et al. 2005; Young et al. 2006; Retting et al. 2009; Komori 2010; Medio et al. 2012). Correspondingly, molecular evolution of candidate developmental genes PAX9 (Phillips et al. 2013) and RUNX2 (Ferraz et al. 2018) suggests that the mechanism of osteoblast differentiation is related to phyllostomid facial length variation. However, the molecular mechanisms in phyllostomid facial skeletal development and evolution have never been directly examined. To assess the pattern of progenitors in phyllostomid craniofacial development and evolution, we examined the CNC and a key mechanism of their proliferation and differentiation—the BMP (bone morphogenetic protein) signaling pathway.

### BMP signaling key regulator of CNC and their derivates

BMP signaling, initially discovered for its role in cartilage and bone development, is important to normal craniofacial development. Targeted mutations of the pathway in CNC, either by ligand removal or receptor defects, alter signaling and result in craniofacial anomalies such as a shortened midface (Kanzler et al. 2000; Matsui and Klingensmith 2014), cleft lip (Zhang et al. 2002; Li et al. 2013), and cleft palate (Wyatt et al. 2010; Saito et al. 2012), and highlight the functional involvement in different aspects of the head (Nie et al. 2006; Wang et al. 2014; Salazar et al. 2016). The canonical BMP signaling pathway has also been shown to regulate facial morphology during evolution. An upstream ligand of the canonical BMP pathway, *Bmp4,* has been implicated in classic models of adaptive radiation—the beak shape of Darwin’s finches (Abzhanov et al. 2004) and the facial shape of African cichlid fish (Albertson et al. 2005). Still, at least 30 upstream factors of the canonical BMP signaling pathway have potential to affect craniofacial development and evolution in mammals, such as *Bmp2, Bmp4, Bmp5, Bmp7,* and *Msx1/2* (Kanzler et al. 2000; Brugmann et al. 2006; Chai and Maxson 2006; Nie et al. 2006; Guenther et al. 2008; Bonilla-Claudio et al. 2012). Further, two key antagonistic regulators of the canonical BMP signaling pathway, Noggin and Gremlin, are required for craniofacial and cartilage development (Merino et al. 1999; Wordinger et al. 2008; Zuniga et al. 2011; Al Chawa et al. 2014; Matsui and Klingensmith 2014).

In an effort to understand the evolution of facial length diversity, we describe, quantify, and compare the tissue-level spatial pattern of cranial neural crest-derived progenitors and the BMP signaling pathway during midfacial development among bat species that vary in facial length. We compare BMP expression at CS18, a stage where we previously have shown proliferation and facial length to be different between species, and at CS17, a stage preceding species-specific variation in facial length (Figure 1). Since numerous ligands and receptors are involved in BMP signaling and the expression of these does not equal BMP activity, we directly investigate *in vivo* BMP signal transduction (i.e. activity) using phospho-SMAD 1/5/8 (9) (pSMAD) immunostaining (Leeuwis et al. 2011; Huycke et al. 2019). Using this approach, we found that the spatial expression pattern of pSMAD underscored facial length at CS18 and, unexpectedly, the presumptive leaf-nose of phyllostomid bats at CS17 and CS18. To evaluate further, we used mouse genetics to manipulate BMP activity during midfacial development. These mouse experiments demonstrated that the BMP-responsive regions of the frontonasal region contributed to morphological variation analogous to aspects of bat face development, particularly in facial elongation and nasal expansion. Phyllostomid bats seem to be particularly susceptible to having these changes contribute to their dazzling array of facial diversity however, it is likely other bat groups have similar frontonasal progenitor modifications by BMP signaling, like the convergent old world leaf-nose bats (Family Hipposideridae, Rhinolophidae, Rhinonycteridae, Megadermatidae).

**Fig (1).**
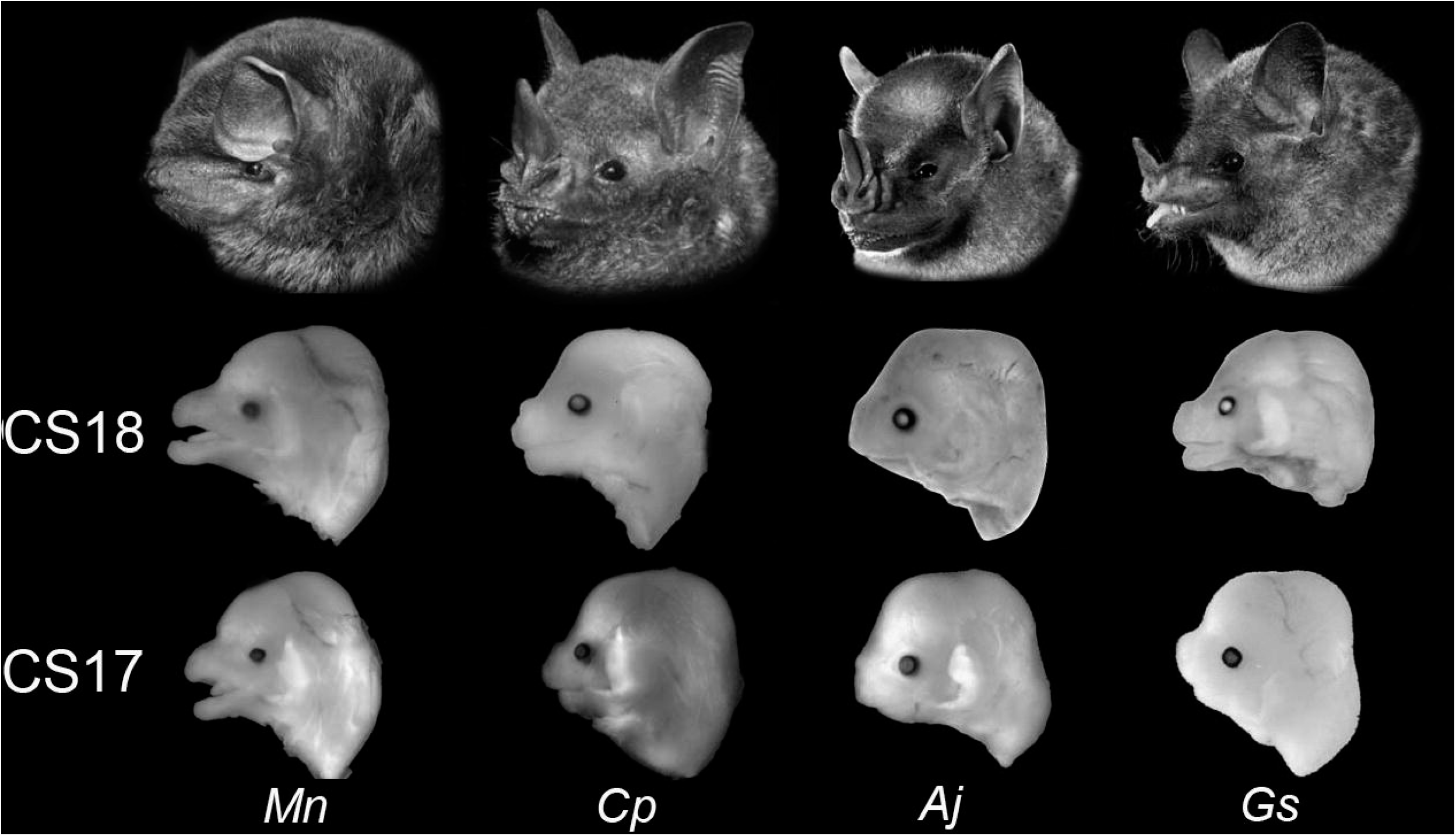
Embryonic cranial shape in bats. Lateral views of the four bat species with different diets, morphology, and embryonic age. Speciesspecific midfacial length morphology emerges during the five day interval between CS17 and CS18 among phyllostomid bats. The long, slender face of outgroup species *M. natalensis* (left-most) is apparent at CS17. In all phyllostomid species, the midfacial length differences are species-specific at CS18. Species shown from left to right: *M. natalensis (Mn), C. perspicillata (Cp), A. jamaicensis (Aj)*, and *G. soricina (Gs)*. Species are scaled to the same cranial length at each stage.

## Results

### Craniofacial variation in bat development

Prior to molecular analysis, surface landmark based geometric morphometric analysis (Supplemental Figure 1) of bat embryo heads in lateral view were used to confirm previously described morphological variation of the craniofacial skeleton (Camacho et al. 2019). It also added new descriptions of soft-tissue development. The first two principle components (Figure 2A) capture the majority of shape variation (68% variation). PC1 (48% variation) describes dorsal growth of the forebrain, anterior growth to the maxilla, dorsal-ventral expansion of the nasals and leaf-nose lancet, growth of the external ear tissue, and rotation of the midbrain. PC2 (20% variation) describes caudal expansion in forebrain, midbrain and hindbrain, anterior expansion of the lancet and lips, elongation of the midface and mandible, and the anterior elongation of the external ear tissue.

**Fig (2).**
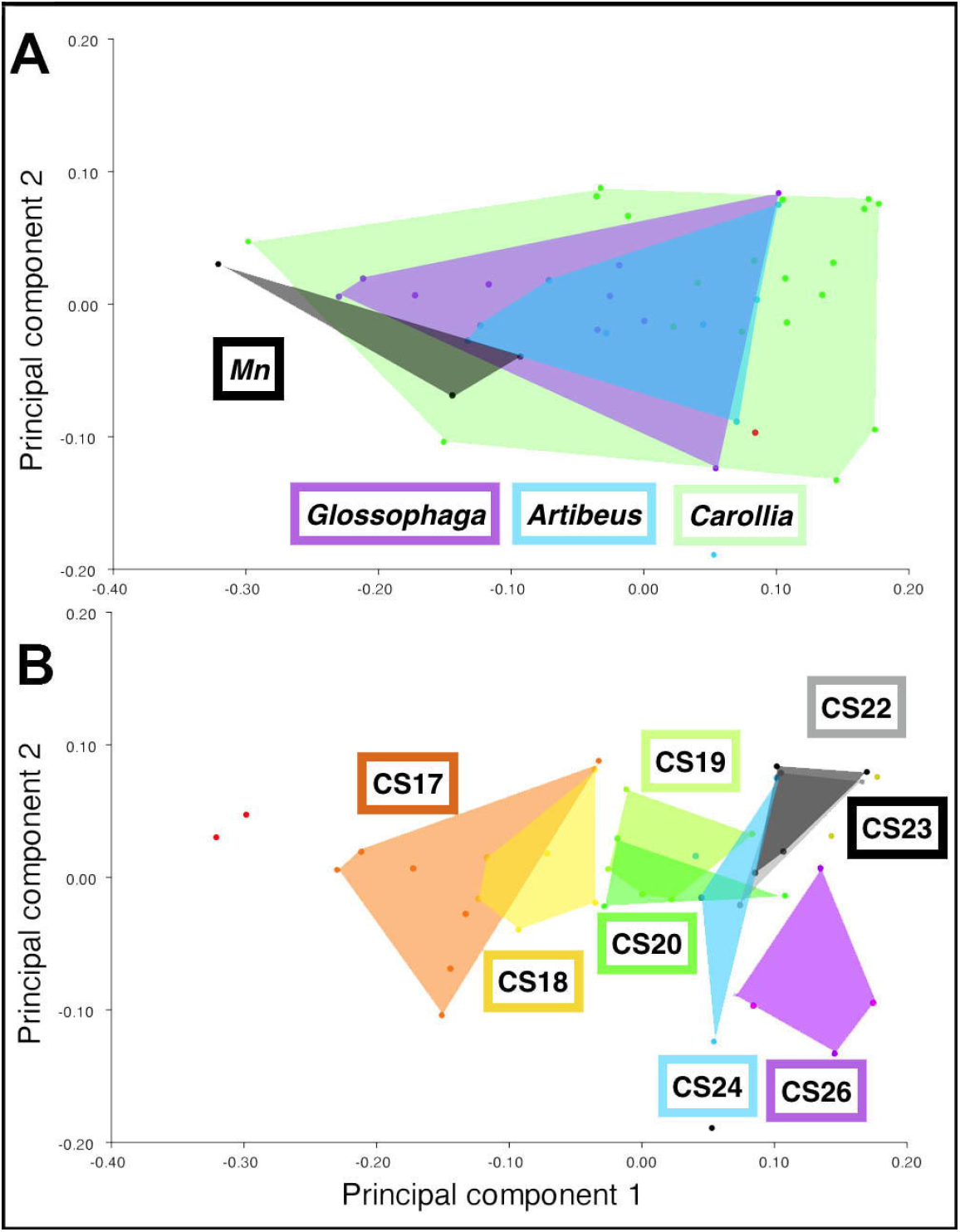
Morphospace plots of craniofacial development. Principal components (PC) analysis defined by the first two PCs of shape-data for *C. perspicillata (Carollia), A. jamaicensis (Artibeus),* and *G. soricina (Glossophaga)* during craniofacial development (CS16-CS24+). Principal component (PC) 1 explains 68% of variation and PC2 explains 48% of variation between species. Data points are color-coded by species as indicated in A. There is broad overlap in morphological changes among bat species (colored-convex hulls) during development. Variation along PC1 and PC2 is more restricted in *Artibeus* compared to *Glossophaga* and *Carollia. M. natalensis (Mn)* occupies a small region of morphospace, possibly related to a more limited sample range (CS16-CS18) (A). The separation along PC1 relates to stage of development, regardless of species. As development progresses, PC1 scores increase. Each developmental stage is grouped as a convex hull. CS19 and CS20 are similar along PCI, but distinct along PC2, and CS22 and CS23 occupy the same region of morphospace (B).

Along the morphospace defined by PC1 and PC2, species within the phyllostomid family are more varied in shape during their development compared to the outgroup, which is restricted to a smaller portion of morphospace (Figure 2A). Distinct clusters are apparent when considering developmental stage (Figure 2B). Morphologically, species at the same stage are more similar to each other at the earlier stages of development (p-adj <0.05). At CS19 and CS20, the morphological variation is similar along PC1 and distinct along PC2. At CS22 and CS23, which is represented only by *C. perspicillata,* the morphologies along PC1 and PC2 occupy the same region of morphospace. At fetal stages, CS24 and CS26, there is a shift toward negative PC2 values.

### Cellular mechanism of craniofacial variation in bats

To understand the differences in facial length between CS17 and CS18, we investigated the progenitors of the midface, the cranial neural crest (CNC), with Pax7 immunohistochemistry. Pax7 is a transcription factor marker for neural crest in mouse, chick, zebrafish, and frog (Basch et al. 2006; Otto et al. 2006; Lacosta et al. 2007; Yoshida et al. 2008; Maczkowiak et al. 2010; Murdoch et al. 2012). At CS17 through CS18, in all bat species, Pax7 is detected in the frontonasal region (Figure 3, 4). It is found in the nasal cartilage, the frontonasal mesenchyme, the undifferentiated mesenchyme of the future olfactory turbinates, the olfactory epithelium, skin, and whisker follicle. It is also found in other non-neural crest cranial regions, like the pituitary gland and brain, eye lens, neural retina, ocular muscles, and eyelids (data not shown). Pax7 is more widely detected at CS17 (Figure 3), and then becomes rostra I ly restricted along the midline in the frontonasal regions by CS18, whereas Pax7 becomes caudally restricted along the lateral frontonasal region (Figure 4). Overall, the differences in the amount and distribution of Pax7-positive cells between species are not statistically significant after correcting for multiple comparisons (Supplemental Figure 3).

**Fig (3).**
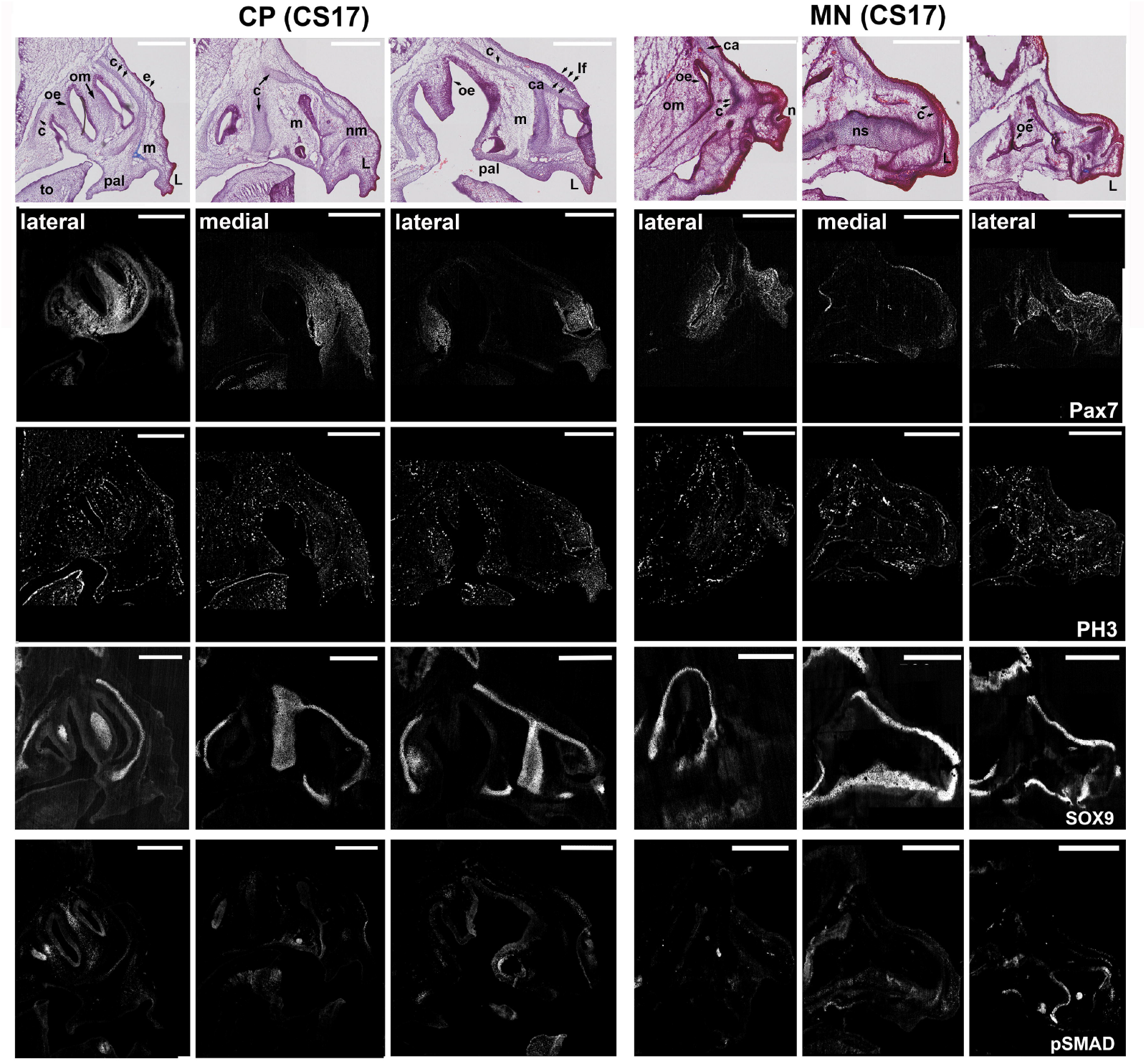
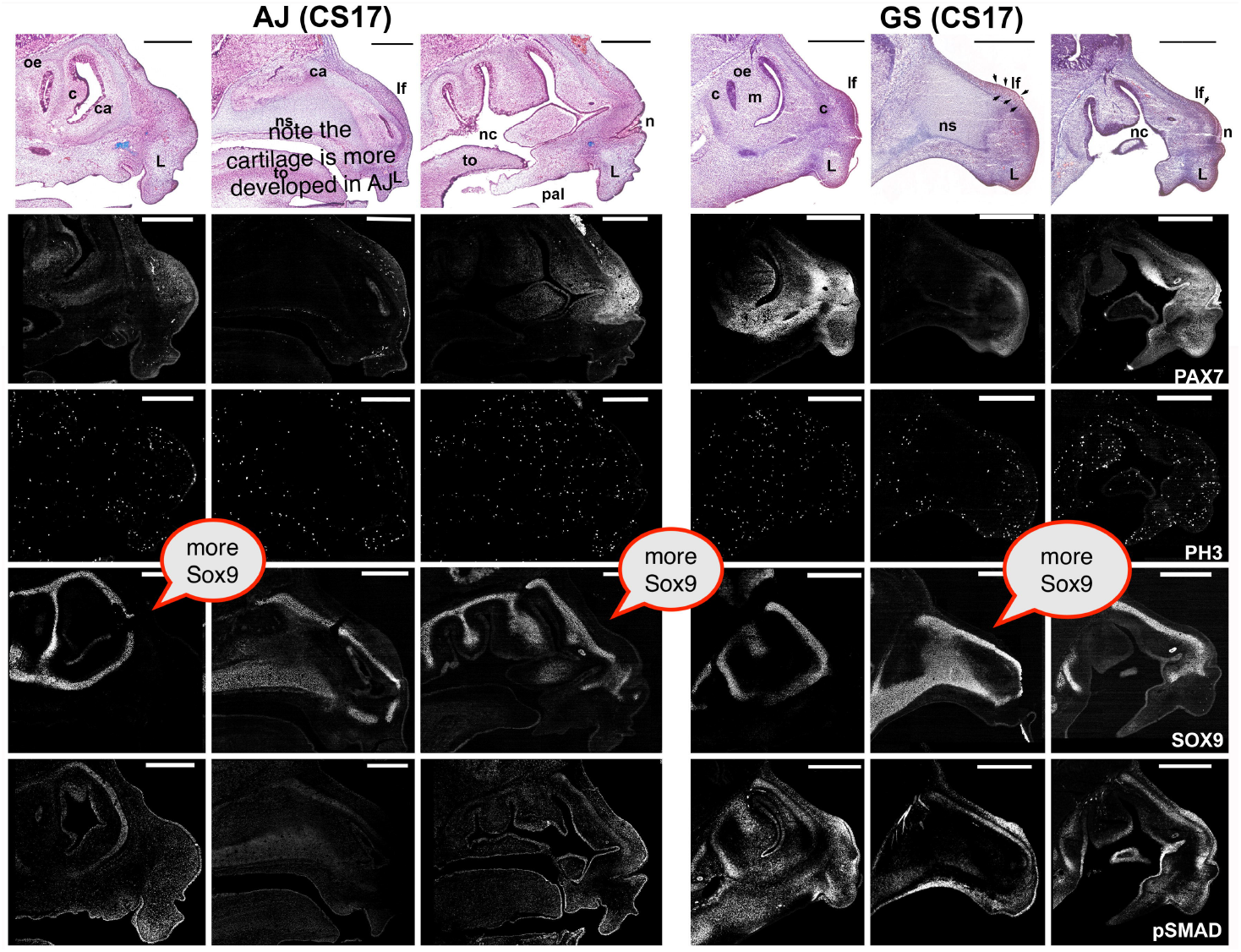
Spatiodistribution of molecular markers Pax7, PH3, Sox9, and pSMAD at CS17. Immunofluorescence targeting the CNC (Pax7), mitotic cells (PH3), cartilage (Sox9), and BMP signaling (pSMAD) in bat embryos are represented by three sagittal sections capturing the medial (middle) and lateral aspects of the midface. Pax7 and PH3 immunostaining are performed on the same section, whereas pSMAD and Sox9 immunostaining are performed on adjacent sections approximately 10 and 20um apart, respectively, from the Pax7;PH3 double-immuno. Histological staining with trichrome (top of panel) match the three different levels. Expression of Pax7 is detected in multiple cranial regions, but is primarily enriched in the skin (epithelium), the upper lip mesenchyme (L), nose mesenchyme (nm), leaf nose (If), and the developing olfactory turbinates (om). In the cartilage (ca), a distinct spatial pattern is observed, with Pax7 expressed in lateral, but not in medial cartilages. PH3-positive mitotic cells are observed throughout the midface, but the medial cartilage has fewer mitotic cells. Sox9 expression is observed in the early developing cartilages of the midface with distinct patterns in the lateral and medial regions among species. pSMAD expression is observed in both Pax7 (undifferentiated mesenchyme, m) and in Sox9 (early cartilage commitment) spatial domains, as well as the developing secondary palate. CP (*Carollia perspicillata*); MN (*Miniopeturus natalensis*); AJ (*Artibeus jamaicensis*); GS (*Glossophaga soricina*). C = condensed mesenchyme; ca = cartilage; e = epithelium; L= lip; If= leaf nose; m = mesenchyme; n= naris; nc= nasal cavity; nm= nasal mesenchyme; ns= nasal septal cartilage; oe = olfactory epithelium; om = olfactory mesenchyme; pal = palate; t= tooth; to = tongue. Scale: 500um

**Fig (4).**
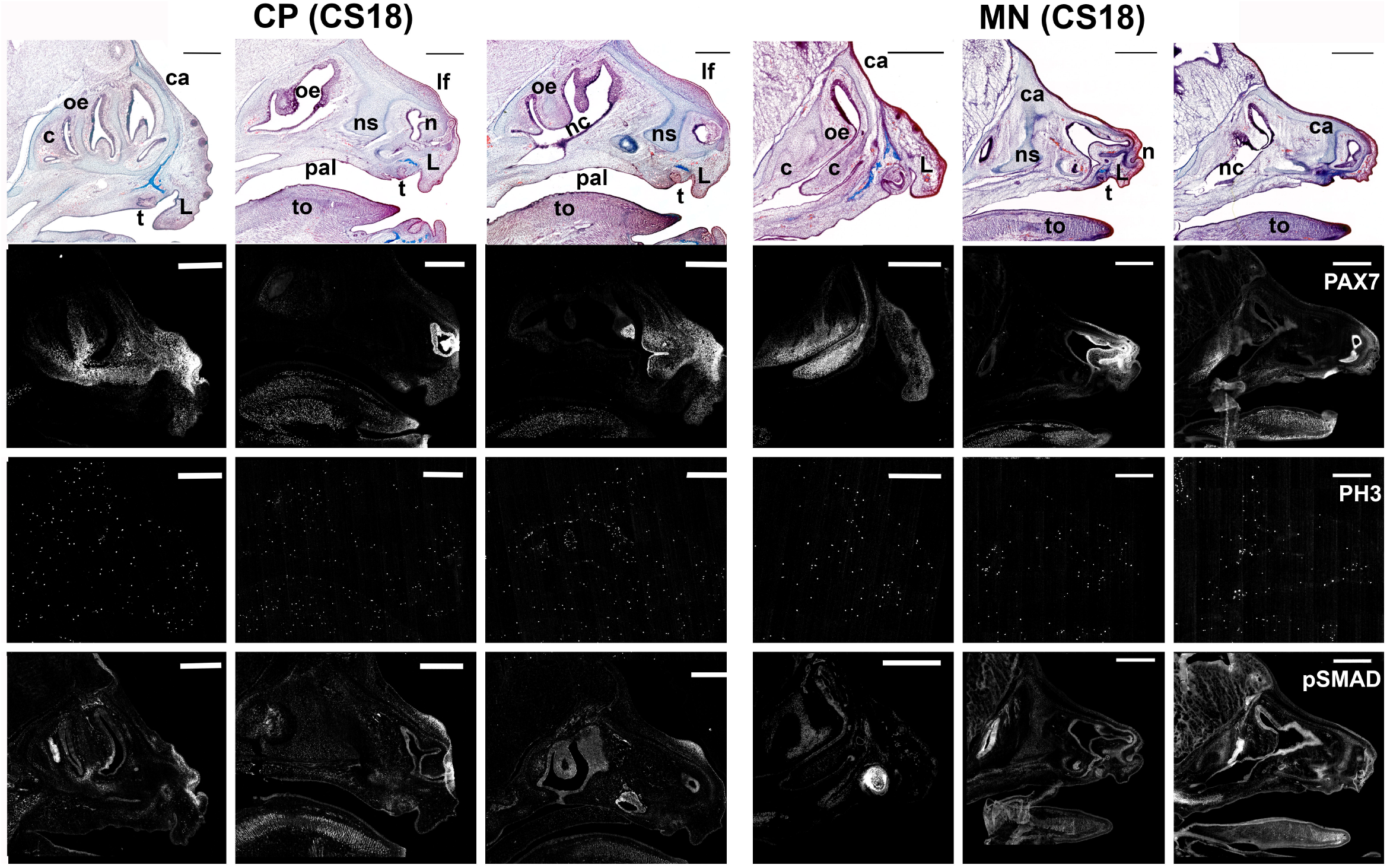
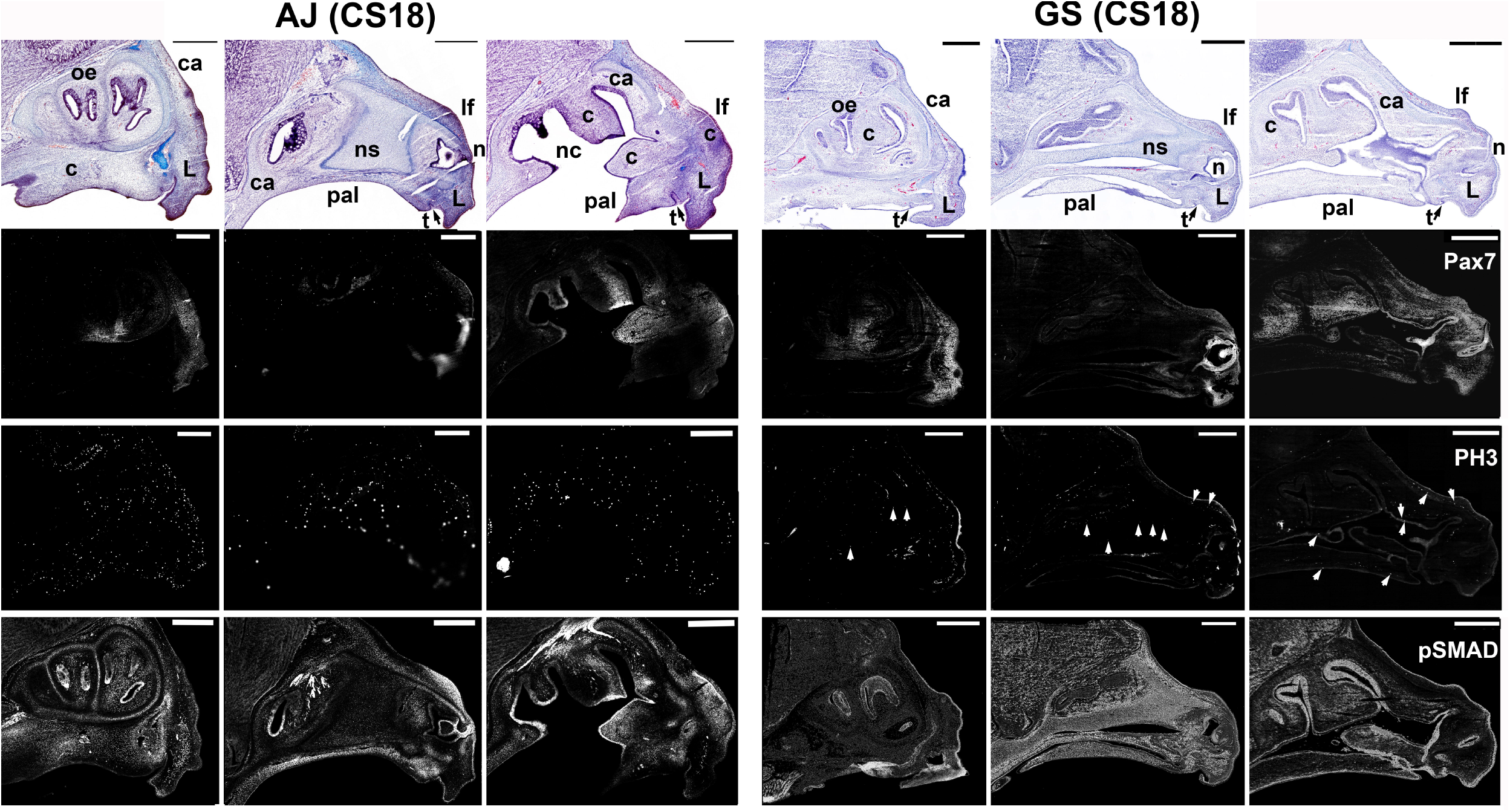
Spatiodistribution of molecular markers Pax7, PH3, and pSMAD at CS18. Immunofluorescence targeting the CNC (Pax7), mitotic cells (PH3), and BMP signaling (pSMAD) in bat embryos at CS18. Top panel shows histological staining with trichrome highlighting cartilage and connective tissue development (blue) and each fluorescence image below match the three sagittal sections capturing the medial (middle) and lateral aspects of the midface. Expression of Pax7 is detected anteriorly in the developing upper lip (L) and posteriorly in the condensing (c) mesenchyme of the olfactory turbinates. It is also observed in the developing leaf-nose (If), tongue (to), and around the naris (n). It is not present in the mature, differentiated cartilage. Based on histology and Pax7 expression, the olfactory turbinates are more mature in *Artibeus jamaicensis* (AJ), which express less Pax7, than other bats at CS18. Laterally (far left), *Carollia perspicillata* (CP) is highly expressed along the cheek tissue and condensed mesenchyme (c) of the developing turbinates. PH3-positive cells are observed in the midface, but are more highly present in AJ and CP than in GS and MN. Arrowheads highlight some PH3-positive cells in GS, which have an excess in non-specific background staining. In all species, pSMAD expression is similarly distributed throughout the midface in the olfactory epithelium (oe), cartilage (ca), condensing mesenchyme (c), palate (pal), tongue (to), nasal septal cartilage (ns), and leaf-nose (If). pSMAD is more highly expressed in the developing palate and the olfactory turbinates of AJ. pSMAD is more highly expressed throughout the nasal septal (ns) cartilage of GS. We note an excess in background staining in our pSMAD immunos for GS. Pax7 and PH3 immunostaining are performed on the same section, whereas pSMAD immunostaining is performed on an adjacent serial section approximately 10um from the Pax7;PH3 double-immuno. CP (*Carolliaperspicillata*); MN (*Miniopeturus natalensis*); AJ (*Artibeus jamaicensis*); GS (*Glossophaga soricina).* C = condensed mesenchyme; ca = cartilage; e = epithelium; L= lip; If= leaf nose; m = mesenchyme; n= naris; nc= nasal cavity; nm= nasal mesenchyme; ns= nasal septal cartilage; oe = olfactory epithelium; om = olfactory mesenchyme; pal = palate; t= tooth; to = tongue. Scale: 500um

The spatial expression of Pax7 and their corresponding histological morphology suggest the Pax7-positive cells were the CNC-derived mesenchyme (ectomesenchyme). Thus, we examined the potential progenitors by evaluating proliferation with molecular marker phosphohistone-H3 (PH3) as previously described (Camacho et al. 2020). We evaluated if facial length may be the result of proliferation differences in ectomesenchyme (Pax7-positive) or the differentiated ectomesenchyme-derived tissue (Pax7-negative). We found that the Pax7-positive ectomesenchyme was highly proliferative compared to surrounding Pax7-negative tissues (Supplemental Fig A3). To understand how the spatial and temporal differences of Pax7 progenitors along the frontonasal region contribute to facial length, we quantified and compared Pax7-positive cells during facial elongation. For each species, the population of Pax7 cells were found to decrease as facial length increased from CS17 to CS18 (Supplemental Figure A4).

The majority of proliferation along the midline of the face is localized to Pax7-positive tissue at the rostral end of the face, in the frontonasal mesenchyme (Figure 3, 4). Other Pax7-positive tissues with relatively elevated proliferation, compared to Pax7-negative tissues, are located at the lateral edges of the face—to the mesenchymal condensations of the developing olfactory turbinates—and are directed along a dorsoventral direction (Figure Supplemental A3). In line with our previous reported results on proliferation, the majority of proliferation directed along the anterior to posterior direction is localized to Pax7-negative tissue along the midline and caudal end of the face (Figure 4, arrows). Anatomically, this relates to chondrocytes of the developing nasal septum and presphenoid. Molecularly, the downregulation of Pax7 relates to the onset of cellular differentiation (Chen et al. 2006).

Our data suggest that cell proliferation and subsequent differentiation to chondrogenic potential, likely within the nasal septal cartilage, contribute to facial length differences between bats. We observed early, but faint, histological evidence of cartilage cells with Masson’s trichrome staining for collagen along the length of the face at CS17 (Figure 3). When we look at the expression of molecular marker Sox9, a transcription factor important to chondrocyte differentiation (Bell et al. 1997; Mori-Akiyama et al. 2003), Sox9 is restricted in its expression to the ectomesenchymal condensation at CS17 in all species (Figure 3). The less elaborate olfactory turbinates of *M. natalensis* expresses less Sox9 (Figure 3A) compared to the olfactory turbinates of phyllostomids. Within phyllostomids, *G. soricina* appears to have elevated expression of Sox9 (Figure 3B), however variation in sectioning angle made direct comparisons of the midline structure development more challenging compared to lateral structures. At CS18, cartilage development is expanded (Figure 4). Since Sox9 is activated through the BMP signaling pathway (Zehentner et al. 1999; Pan et al. 2008), we focused next on evaluating BMP signaling in craniofacial development.

### Nuclear pSMAD is broadly expressed in craniofacial development

In the nucleus, phospho-SMADs of the BMP pathway regulate gene expression controlling cell proliferation and differentiation of pre-cartilage and pre-bone cells (Wrighton et al. 2009). We evaluated BMP-responsive signaling activity with nuclear phospho-SMAD 1/5/8(9) immunohistochemistry (Retting et al. 2009; Wrighton et al. 2009; Feng et al. 2015) in bat embryos. At CS17, BMP is active in the developing nasal septal cartilage, presphenoid, palate, and frontonasal mesenchyme (Figure 3), as well as some regions of the brain (data not shown). At CS18, the expression domain of BMP was expanded to the growing nasal septum, the growing edges of the olfactory nerves and epithelium, the leaf-nose, palate, canine tooth bud (Figure 4), the sphenoid sinus, the body of the tongue, the choroid plexus, and resting cartilage cells of the cranial base (Supplemental Figure 5). The expression of the BMP signaling in the midfacial regions is associated with cartilage and bone growth, which have been shown to affect facial length development (Pavlov et al. 2003; Dudas et al. 2004; Parsons et al. 2015; Kaucka et al. 2017).

Molecular comparisons of BMP signaling were investigated further in the midface at CS17 and CS18. At CS17, no differences are found in pSMAD expression among species (Figure 5). At CS18, BMP expression showed elevated expression in the long-face nectar bat *Glossophaga soricina* when compared to the shorter faced model bat *Carollia perspicillata* (Figure 5). In contrast, BMP activity is decreased in the short-face fruit bat *Artibeus jamaicensis* when compared to the slightly longer faced model bat *C. perspicillata* (Figure 5). Overall, the changes in pSMAD expression relate to the developing nasal septal cartilage and to a lesser extent the palate (Figure 6).

**Fig (5).**
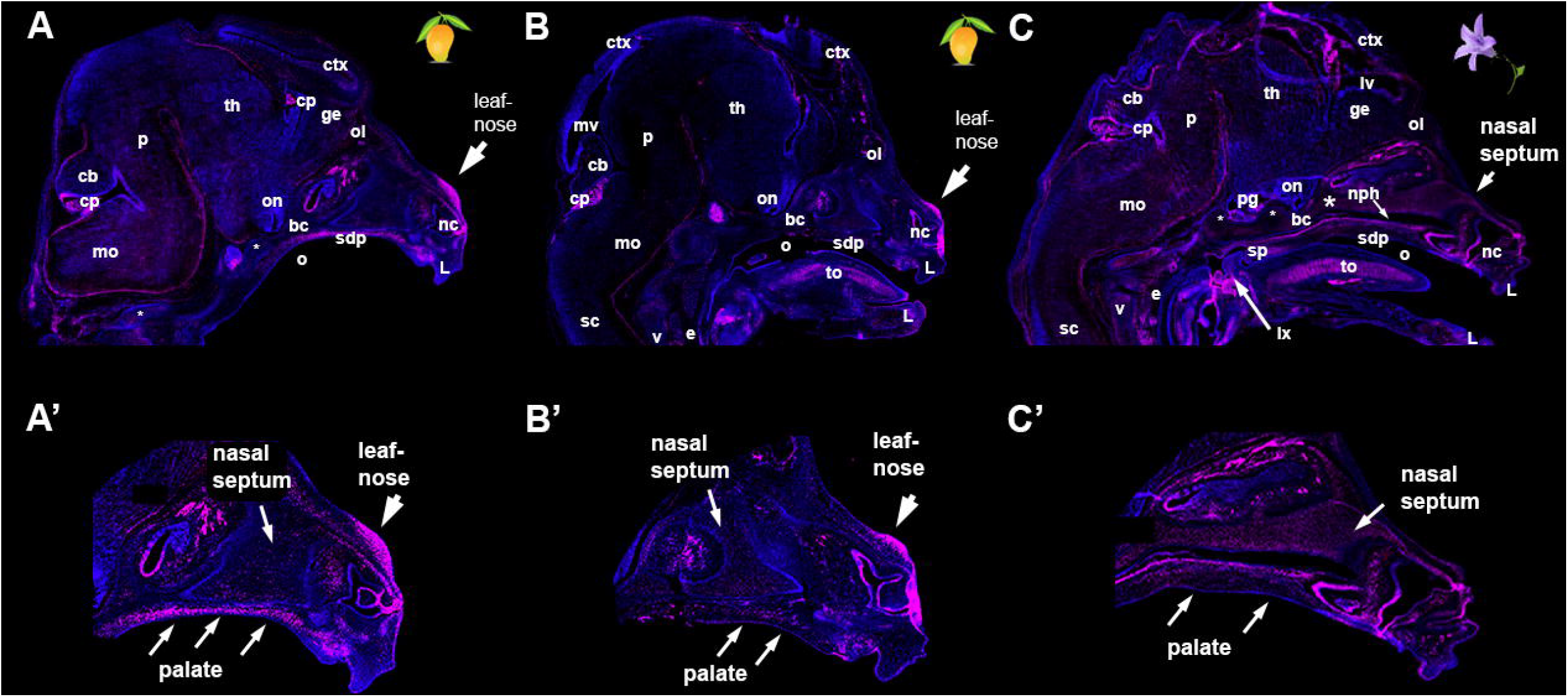
BMP signaling domains related to embryonic facial morphology. Relative expression changes in overall BMP-signaling pathway (pSMAD 1/5/9 immunohistochemistry). These representative parasagittal sections highlight species-specific differences in pSMAD molecular expression (magenta) and related cranial morphology (DAPI, blue) in the fruit-eating (A, B) and nectareating bats (C). The pSMAD expression along the basicranium relates to facial length morphology: pSMAD is less expressed throughout the facial cartilages *of A. jamaicensis* compared to *C. perspicillata* and pSMAD is more highly expressed throughout the nasal septal cartilage and basicranium (bc) of *G. soricina* (C, C’) compared to *A. jamaicensis* (A’) and *C. perspicillata* (B’). The elevated pSMAD in the developing secondary palate (sdp) *of A. jamaicensis* (A, A’) might relate to the width of the palate. Cranial features relating to elevated pSMAD expression are the leaf-nose, nasal epithelium, choroid plexus (cp), tongue (to), and larynx (lx), bc= basicranium; cb= cerebellum; cp = choroid plexus; ctx = cortex, ge = ganglionic eminence; L = lip; lx = larynx; mo = medulla oblongata; mv = mesencephalic ventricle; nc = nasal cavity; nph = nasopharynx; o = oropharynx; ol = olfactory lobe; on= optic recess; p= pons; pg = pituitary gland; sc = spinal cord; sdp = secondary palate; sp = soft palate; th = thalamus; to = tongue; vc = vertebrae cartilage.

**Fig (6).**
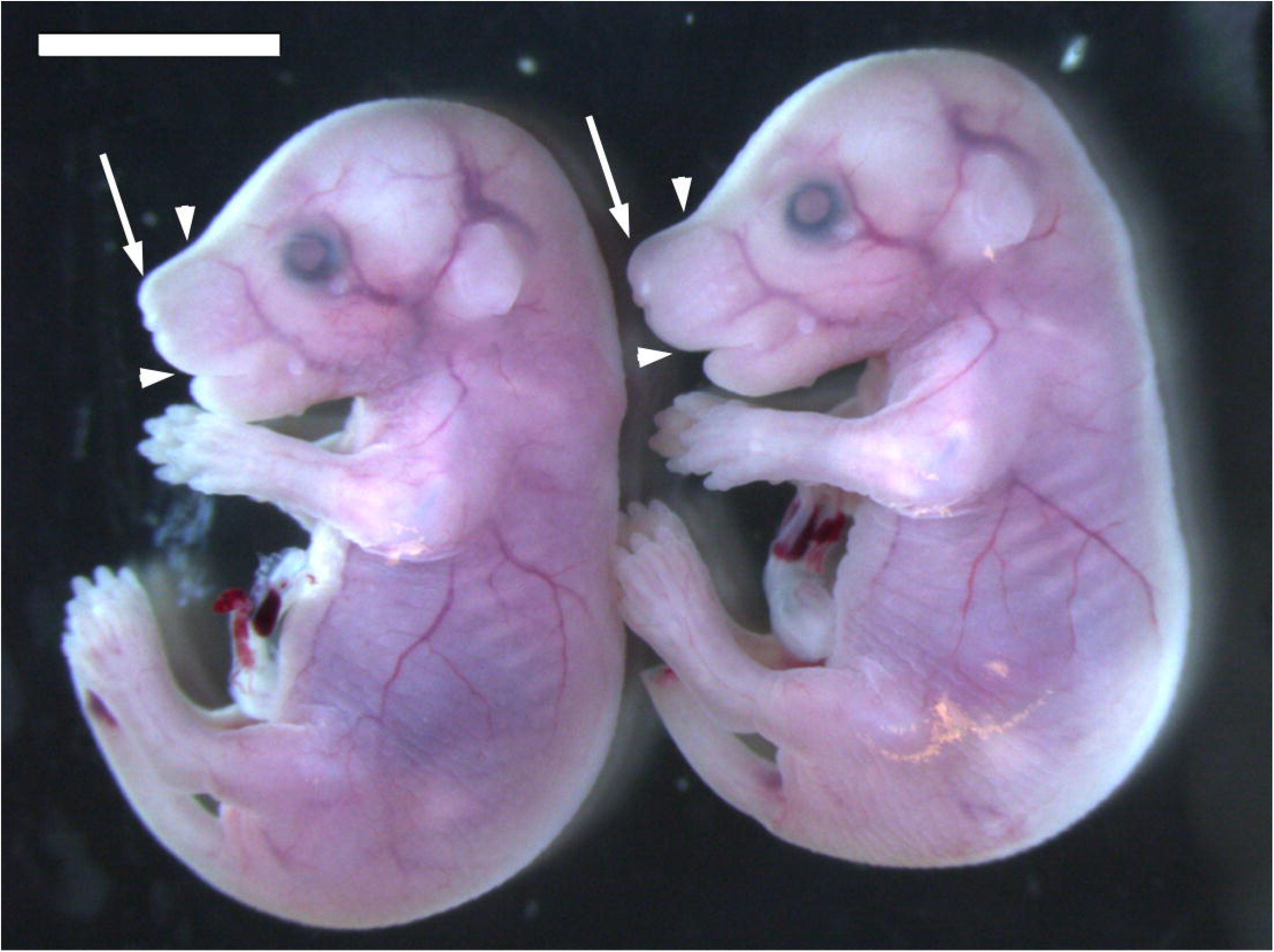
Mouse mutants & FL→features highlighted. Variation in BMP signaling at E12.5 affects facial morphology in E16.5 mouse embryos. Relative to control littermates (left), the *Wnt1* mutant mice (right) have enlarged facial features. The facial length is longer and extends beyond the lower jaw (arrowheads). The rhinarium, or nose, is larger in mutants (arrow).

### BMP mouse mutants mimic craniofacial variation in bats

To evaluate the connection of BMP signaling differences at CS18 to morphological variation, we overexpressed BMP during mouse craniofacial development at ~E12.5/E13.0, which roughly relate to ~CS17/CS18 (Sears et al. 2006; Cretekos et al. 2008; Hockman et al. 2009), and then assessed how the BMP pathway affected mouse facial morphology at E16.5. We used a genetic approach to alter BMP ligand signaling from two main cellular sources: the cranial neural crest (CNC) and mesoderm. Using neural crest *Wnt1*-cre and mesoderm *Twist2*-cre drivers, we conditionally deleted BMP antagonists Noggin and Gremlin at E12.5, a critical period in midfacial morphogenesis associated with skeletal and cartilage tissue development. While not examined here, our lab has recently published that these mice have increased BMP signal activity in various tissues derived from neural crest and mesoderm (Huycke et al. 2019). Both *Wnt1*-cre and *Twist2*-cre mice exhibit noticeably longer snouts compared to wildtype littermates (Figure 7). No gross morphological defects were observed in the external surface morphology, but a noticeable enlargement of the midface was observed. We further evaluated morphology with surface landmark based geometric morphometrics and principal components analysis (PCA), as was done in bats.

**Fig (7).**
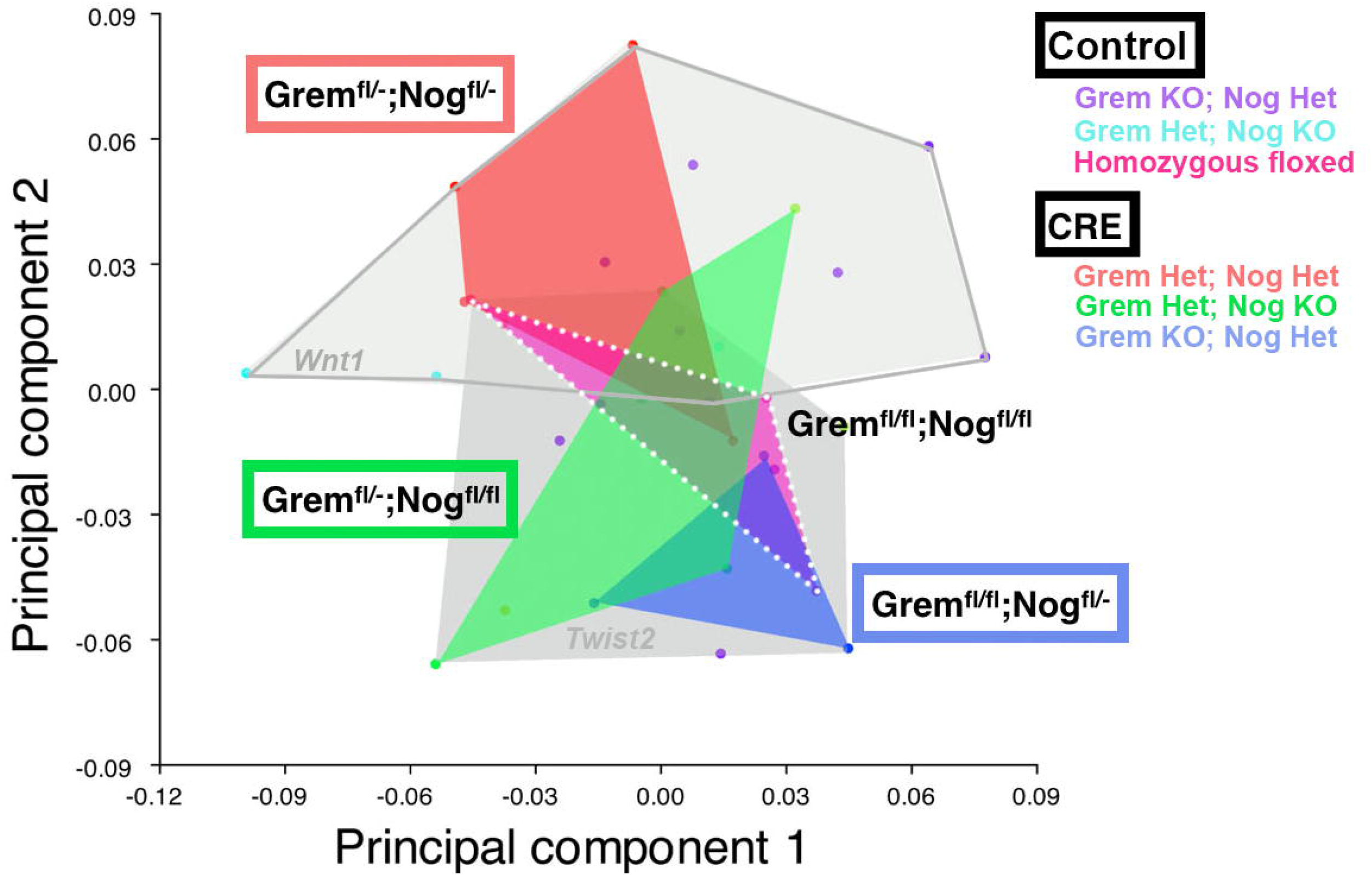
Mouse PCA. Principal components analysis (PCA) defined by the first two PCs of shape-data in mice with deleted Bmp antagonists Noggin and Gremlin driven by *Wnt1-cre* or *Twist2-cre.* PCA captures 45% of cranial variation. PC1 (23% variation) and PC2 (22% variation) cluster mice by cre driver (*Wnt1* or *Twist2)* and by genotype (colored convex hulls). *Wnt1* embryos (n= 2 litters) occupy the upper portion of morphospace and are outlined in grey. *Dermo1* embryos (n=2 litters) occupy the lower region of morphospace (grey convex hull). Both *Wnt1* and *Twist2* are represented by different genotypes: Gremlin homozygous (fl/fl) or heterozygous (fl/-) and Noggin homozygous (fl/fl) or heterozygous (fl/-). Mice homozygous for both Gremlin (fl/fl) and Noggin (fl/fl) did not survive to E16.5. Morphology of mutants are compared to floxed controls (pink convex hull, white dotted outline) at E16.5.

PCA at E16.5 captured 45% of shape variation within the first two principal components (Figure 8). PC1 (23% variation) describes anterior growth to the snout, expansion of the nasal tissue, upward growth of the external ear tissue, and ventral rotation of the midbrain. PC2 (22% variation) describes dorsal expansion in forebrain and midbrain growth, variation in the size of the eye, anterior expansion of the nose, elongation of the midface and mandible, and anterior elongation of the external ear tissue. Along the morphospace defined by PC1 and PC2, mice cluster within genotype (Figure 8). The *Wnt1*- and *Twist2*-cre homozygous control mice (Noggin^flx/flx^; Gremlin^flx/flx^) make up the pink convex hull at the center of morphospace. *Wnt1*-ere floxed mice (increased BMP activity) move away from the control mice and occupy the upper grey convex hull. An increase in BMP activity affected morphology in *Twist2*-cre floxed mice, which occupy the lower grey convex hull in morphospace. Both grey mutant clusters occupy a wider range of morphospace than controls. All mutant mice have similar variation along PC1 and PC2, but *Wnt1*-ere mice have more extreme PC2-positive scores, which relate to their enlarged midface.

**Fig (8).**
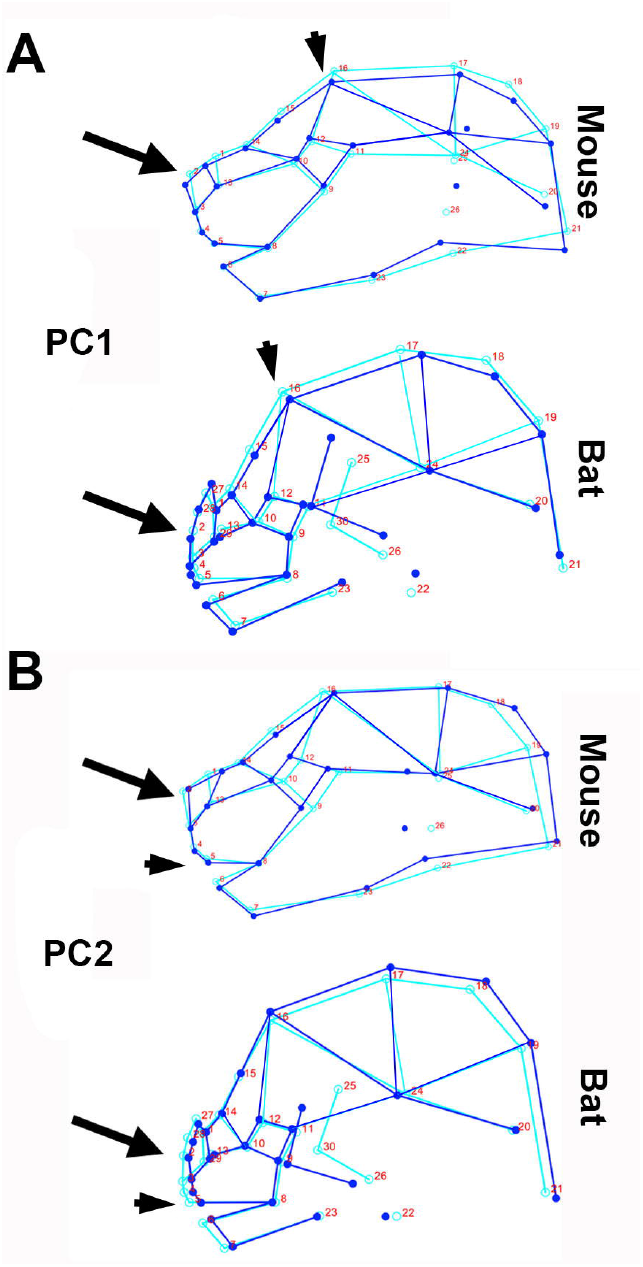
PC1 and PC2 variation in mice resemble bat. Wireframes illustrate major changes across each of the first two principal component (PC) axes. PC1 and PC2 of Bmp mouse mutants are shown alongside PC1 and PC2 of bat cranial development. The dark blue lines represent the mean shape and the light blue lines represent the variation. The arrowheads emphasize the midfacial enlargement in Bmp mouse mutants that resemble midfacial development in bats. The arrow highlights the mouse mutant expansion of the rhinarium that matches the leaf-nose development in bats.

In Wnt1-cre, the ratio of facial length to cranial length is 5% higher than controls, but in *Twist2*-cre, the ratio of facial length to cranial length does not differ from the controls (Figure 7). In addition to a localized increase in midfacial length, the dorsal nasal tissue was expanded in *Wnt1*-cre animals. These phenotypic changes described in mice with altered BMP activity resemble the changes during bat facial development at the morphological level (Figure 9).

**Fig (9).**
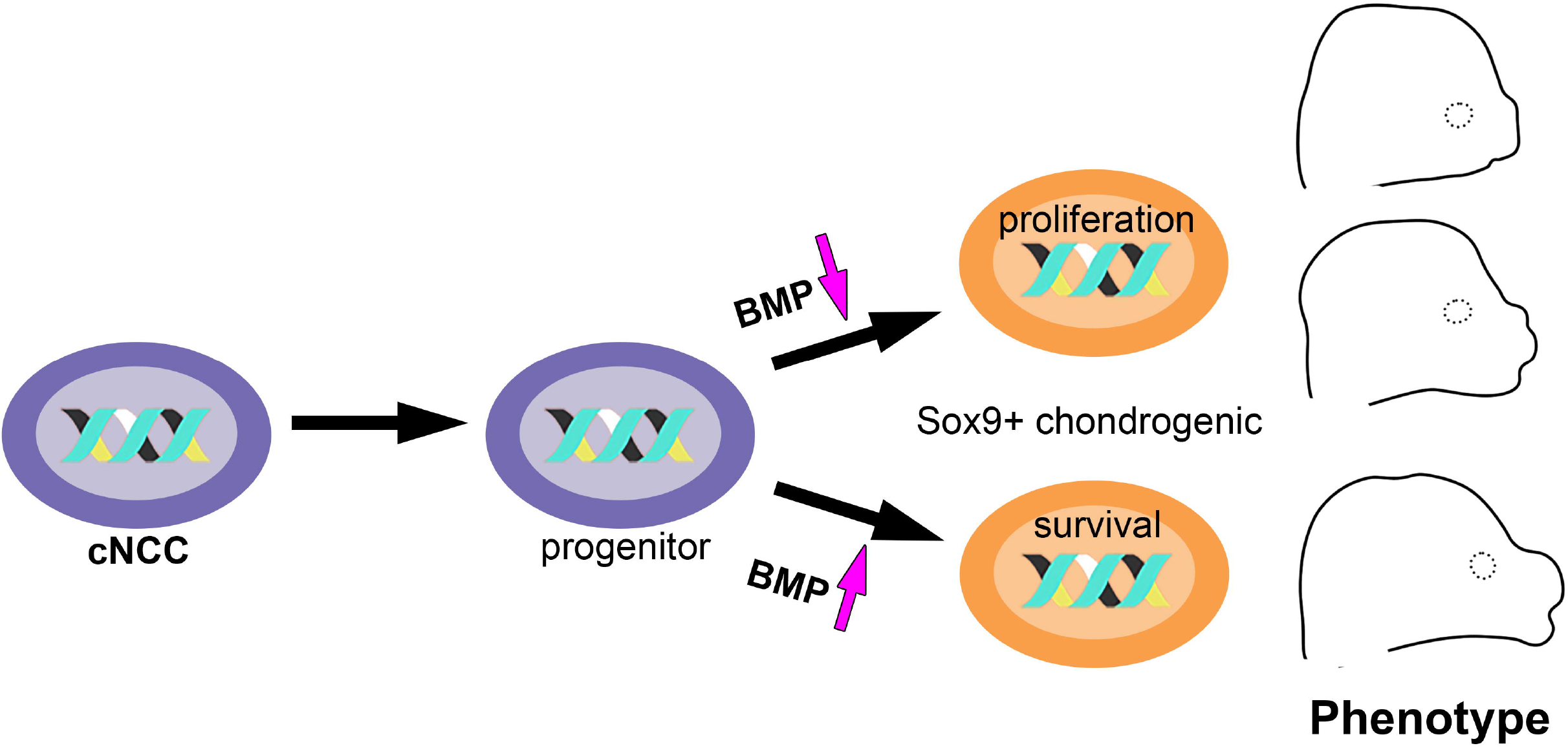
Proposed model where BMP regulates genes involved in proliferation and differentiation of progenitors. Model of Bmp signaling implicated in skeletal cell differentiation and facial length phenotype. Bmp signaling and its modulation are involved in craniofacial skeletal development. Neural crest derived mesenchymal progenitors are specified to chondro-progenitor cells (Sox9+) prior to cartilage maturation. When Bmp signaling is low, chondrocyte proliferation is promoted, resulting in short faces. When the BMP is elevated, cell proliferation is suppressed, resulting in an enlarged midface. Bmp signaling in the Sox9+ chondrocyte lineage affects the size of cartilage by suppressing or promoting cell proliferation. Bmp signaling and its modulation are involved in craniofacial skeletal development. Neural crest derived mesenchymal progenitors are specified to chondro-progenitor cells (Sox9+) prior to cartilage maturation. When Bmp signaling is low, chondrocyte proliferation is promoted, resulting in short faces. When the BMP is elevated, cell proliferation is suppressed, resulting in an enlarged midface. Bmp signaling in the Sox9+ chondrocyte lineage affects the size of cartilage by suppressing or promoting cell proliferation. Our proposed model implicates Bmp signaling in skeletal cell differentiation in the cartilage lineage (Sox9+) as one important aspect underlying facial length phenotype through direct regulation of cell behavior.

## Discussion

Among phyllostomids, cranial morphology primarily differs along the relative length of the face (Monteiro and Nogueira 2011; Dumont et al. 2014; Camacho et al. 2019; Hedrick et al. 2019). We sought to understand the developmental genetic mechanism underlying their adaptive variation in facial length. We previously attributed the morphological changes of facial length differences to a form of heterochrony called peramorphosis (Camacho et al. 2019) and also discovered the heterochronic signature at the cellular (Camacho et al. 2020), which served as our starting point to investigate molecular underpinnings driving differential growth of the face. Here, we broadly examined the early outgrowth of the midface, a conserved mammalian process defined by the migration of cranial neural crest into the facial prominences, where they subsequently aggregate, condense, and differentiate into cartilage and bone progenitors. Key to this process is the BMP signaling pathway which specifies and regulates the cell fate of mitoticaIly active progenitors (Bhatt et al. 2013).

We hypothesized that the balance between proliferation and differentiation of the neural crest derived facial structures via BMP signaling might underlie the two models of peramorphosis. Since we know proliferation is elevated in short-faced bats and lower in long-faced bats (Camacho et al. 2020) during an important stage of skeletal development (CS18), we thought pre-cartilage and/or pre-bone progenitors, observed at the histological level as mesenchymal condensations expressing Pax7, might be contributing to these differences. However, our results on the initial progenitor pool condensations of the midface at CS17 were similar among species (supplemental Pax7 quant). In fact, proliferation and BMP signaling were also similar at CS17 (Fig). In contrast, there was a slight difference in the distribution of pre-cartilage progenitors at CS17 based on SOX9 expression (Fig 3). This led us to consider that proliferation differences might instead be regulated by BMP signaling within the cartilage or bone progenitors at CS18.

In short-faced bats, facial structures might mature more quickly, effectively terminating the growth of the midface. This is based on detailed studies in both birds and mice and was first proposed by (Usui and Tokita 2018). For example, if proliferation is experimentally accelerated in chick embryos, *Runx2* maturation signal to bone becomes expressed prematurely and the size of the facial skeleton is reduced (Hall et al. 2014). Likewise, in quail-duck chimeras, the smaller heads of quail naturally express more *Runx2* compared to the long-faced duck (Eames and Schneider 2008; Merrill et al. 2008). In mice, if proliferation is increased in the cartilage progenitors through removal of a cartilage specific negative regulator of chondrocyte proliferation *(Pten),* then premature mineralization and fusion of the anterior cranial base occurs, thereby causing shorter facial length (Ford-Hutchinson et al. 2007). While we did not directly examine the molecular markers of mature cartilage and bone cells (i.e. *Col2a1* and *Osx*), we do observe more cartilage maturation in *A. jamaicensis* at CS18 (Fig 4B) with traditional cartilage staining methods (trichrome histology). It is possible that the elevated proliferation in short-faced species at CS18 results in early maturity of cartilage through accelerated expression of molecular marker *Sox9* and/or early differentiation of bone cells by accelerated expression of *Runx2,* which terminates growth of the midface. Histologically, we do not see bone cells, consistent with the lack of bone ossification of the midface at CS18.

In long-faced bats, we hypothesized that facial structures mature slowly, essentially allowing the facial structures more time to enlarge before terminal differentiation into mature cartilage and/or bone. While this hasn’t been observed in mouse genetic studies, we do know that in quail-duck chimeras, stage-specific alteration in the cell cycle of proliferative cells can led to an increase in facial length (Woronowicz and Schneider 2019). However, in mouse experiments, it was previously shown that a disruption to terminal differentiation of Meckel’s cartilage results in an increase in jaw length (Wang et al. 2013). Long-faces as a result of a delay in terminal differentiation is then a conservative interpretation to low levels of proliferation observed in our data (Fig).

A possible way to produce heterochronic phenotypes is to change the timing of gene expression (Pham et al. 2017). Through careful genetic studies, members of the canonical BMP pathway, such as *Bmp2, Bmp4* and *Bmp7,* have been shown to underlie facial length evolution by affecting either proliferation and/or differentiation (Abzhanov et al. 2004; Wu et al. 2004; Albertson et al. 2005; Okamoto et al. 2006; Parsons and Albertson 2009; Song et al. 2009; Yoon and P 2019). We assayed BMP signal activity for species-specific differences in bats. We observed differential expression of BMP among species with different facial length at CS18 (Figure 4), but not CS17.

In the truncated and wide face *of A.jamaicensis,* elevated pSMAD localized to the palate (Figure). In mice, by way of comparison, the BMP signaling protein *Bmp4* is expressed in the palatal mesenchyme and required for cellular proliferation (Zhang et al. 2002). Unlike mouse mutants, enhanced BMP activity in *A. jamaicensis* does not result in cleft palate along the mediolateral axis (width) of the palate, suggesting BMP in *A. jamaicensis* might uniquely contribute to the widening of the palate (Sorensen et al. 2014).

In the long-faced *G. soricina,* a nectar bat, we noted a localized increase in BMP activity within the nasal septum and anterior cranial base cartilage (Figure C) compared to other species (Figure A, B). In contrast, the short-faced *A. jamaicensis* presented with less pSMAD activity in the facial cartilages compared to *C. perspicillata* (Figure 6). The observed relationship of BMP signal activity in the midline cartilage and facial length fits with what we know about chondrogenesis, where the removal of pSMAD1/5/8(9) in the cartilage lineage leads to smaller cartilage elements (Retting et al. 2009), and the misexpression of BMP ligands *Bmp2* and *Bmp4,* which signal through pSMAD, promote cartilage formation (Duprez et al. 1996; Bonilla-Claudio et al. 2012; Medio et al. 2012).

Possibly, BMP activity in nectar bats maintains the chondrogenic competence, allowing for extended growth of the midface, whereas the decrease in BMP activity in fruit bats promotes proliferation and entry into early terminal differentiation, resulting in a shorter face (Figure 10). These significant changes in BMP signaling during midfacial growth among the various bat species demonstrate that BMP mediated proliferation as early as CS18 underscores differential outgrowth of the midface. Since much is unknown regarding the evolution of long faces in mammals, we were motivated to replicate the stage-specific expression pattern in nectar bats (elevated BMP signaling) with mouse genetics. To increase BMP signaling, two major BMP signaling modulators, Noggin (He et al. 2010) and Gremlin (Merino et al. 1999; Zuniga et al. 2011), which are expressed along the anterior-posterior midfacial cartilage of the head (Fig), were removed. Overexpression of BMP in mice at E12.5 led to elongated faces as well as a dorsal expansion in the nose tip (Figure 7), regions defined by BMP expression at CS18 (Figure 4, 6). In that respect, when BMP signaling in the CNC-lineage in mice is elevated through the removal of Noggin (Matsui and Klingensmith 2014) or *Bmp4* overexpression (Bonilla-Claudio et al. 2012) at E12.5, facial length and internal nasal cartilages were expanded. These previous studies in mice also presented with an increase in the size of the nose tip (Supplemental Figure 6), which were disregarded in the original mutant descriptions (Bonilla-Claudio et al. 2012; Matsui and Klingensmith 2014). Overall, enhanced BMP signaling in mice led to altered craniofacial development that resembles craniofacial variation in bats (Figure 9).

Intensifying BMP signaling in the CNC is likely necessary for the elaboration of tissues during morphogenesis. The leaf-nose, a novel facial structure in phyllostomids, appears at CS17 (Cretekos et al. 2005), in the dorsal frontonasal mesenchyme (Figure 3). Species-specific differences in the leaf-nose morphology were not apparent at either CS17 or CS18; likewise, we did not see differences in the expression of BMP activity in the developing leaf-nose primordia (Figure 6). In our mice, the removal of BMP antagonism might allow BMP expression to be expanded dorsally, affecting the outgrowth of the dorsal structures. In zebrafish, as a point of comparison, dorsal expression of Gremlin in the face restricts BMP to the ventral aspect of the face (Zuniga et al. 2011). By decreasing the dorsal domain of Gremlin, BMP expression became expanded to the dorsal aspect of the face and expanded dorsal cartilage structures (Zuniga et al. 2011). Our results on the unique trait of phyllostomid bats, then, provide an anatomical view of how ligand modulation of BMP signaling in the frontonasal region at a key period of facial development can mimic multiple adaptive and innovative traits observed in nature.

## Conclusion

Comparative embryology of the face between closely related species provides an opportunity to examine how variation arises in the formation of facial structures. These differences highlight possible manners by which evolution has shaped species-specific craniofacial adaptations. Our results suggest BMP signaling controls heterochronic growth of developmentally flexible traits, potentially promoting enhanced proliferation and differentiation. Our findings led us to propose that BMP signaling in CNC promotes the persistence of progenitor cells and the timing of differentiation through targets of BMP signaling, like *Runx2* or *Col2a1*, which underscores peramorphosis in the elaboration of morphological traits in bat facial evolution (Figure 10). More work is still needed to elucidate the role of BMP signaling in bat facial developmental mechanisms, such as cell proliferation and differentiation toward chondrogenic or osteogenic cell fates (Bhatt et al. 2013). Several BMPs have high affinity for Noggin, including BMP2, 4, 6, and 7 (He et al. 2010; Matsui and Klingensmith 2014), any of which might be contributing to the activity of BMP signaling differences among bat species. The connection between the development of bats and the developmental genetics in mice shows how a cellspecific modification in a single pathway can mimic multiple evolutionary traits in bats and potentially underlies how the group has been able to diversify the face so rapidly. Further, morphological transformations by CNC-mediated gene expression involving BMP signaling may also underlie diverse craniofacial phenotypes and novel fleshy facial appendages arising from the frontonasal prominence in mammals.

## Experimental Procedures

### Embryonic tissues from bats

*Carollia perspicillata, Artibeus jamaicensis,* and *Glossophaga soricina* were wild caught from Trindad as previously described in Camacho et al. 2020 under a wildlife collecting permit. Specimens acquired are shown in supplemental Table X. At least three cranial samples at CS17 and CS18 were used for histological and immunostaining.

### Stereomicroscope imaging and biometrics

Embryonic bat heads and embryonic E16.5 mouse heads were submerged in chilled lx PBS and imaged using a stereo microscope with brightfield illumination (Leica M165 FC) in standard anatomical views, which took 5 minutes per embryo. Measurements of the embryo head were obtained with FIJI (Schindelin et al. 2012). For midfacial measurements, the dorsal view of the embryo head was used to capture length of the snout and the cranium. We define facial length as the distance from the nasal tip to the anterior eye and cranial length (CL) as the distance from the nasal tip to caudal head.

### Geometric morphometrics

2D surface landmarks were chosen to capture morphological changes in the frontonasal and maxillary prominences that give rise to the face. Bat embryos in lateral view spanning CS16-CS24 were landmarked in ImageJv1.52e (Schindelin et al. 2012) using the PointPicker plugin following a standard embryo craniofacial landmark set (Percival et al. 2014). In addition, we add additional landmarks to capture bat-specific leaf-nose development (Supplemental Figure 1). The raw 2D coordinates (X,Y) for 30 lateral landmarks were exported from ImageJ as .tps files and analyzed in MorphoJ (Klingenberg 2011). A generalized Procrustes fit was used to create a best fit for landmarks. Similarly, lateral images of four background controls, two Wnt1-Cre; Nog^fx/fx^; Grem^fx/-^, three Dermo1-cre; Nog^fx/fx^; Gre^fx/-^ and three Dermo1-cre; Nog^fx/-^; Gre^fx/fx^ from E16.5 mouse embryos were landmarked in ImageJ with the same landmark set used in bats.

### Histology of sections

To examine cellular morphology of the developing head, 10um serially-cryosectioned bat embryo cranial tissue in sagittal orientation, stored at −80C, were dried to room temperature and fixed at room temperature 10% neutral buffered formalin for 1 hour, followed by an overnight mordant in Bouin’s fixative overnight at room temperature. Afterwards, slides were stained with Masson’s Trichrome (Polysciences) to visualize cell nuclei, muscle, cytoplasm, erythrocytes, and connective tissue.

### Section immunostaining

Bat embryo tissue at CS17 and CS18 had been previously serially-sectioned into 10 x 10um-sagittal orientation (Camacho et al. 2020) and stored at −80C. Three sets of frozen sagittal sections per embryo were immunostained for comparative analysis of facial development. For antigen unmasking, the slides were steamed in pre-warmed 1mM sodium citrate buffer with 0.05%Tween20, pH6.0 for 30 min. Double immunofluoresence for Pax7 (1:250, DSHB) and PH3 (1:1000, Millipore) was performed to detect CNC and proliferating cells with primary antibodies, which were subsequently identified with Alexa Fluor-conjugated secondary antibodies (1:500, Jacksonlmmuno). Immunostaining for Sox9 (1:250, Abcam) was done to detect differentiating cells, which were subsequently identified with Alexa Fluor-conjugated secondary antibodies (1:500, Jacksonlmmuno). Immunostaining for BMP signaling (pSMAD1/5/9,1:500, Cell Signaling) was done, followed with a biotinylated secondary antibody (1:300, Jaksonlmmuno), amplified by a streptavidin-horseradish peroxidase conjugate (1:300) and detected with TSA tyramid-Cy3 (1:50, PerkinElmer). All sections were counterstained with DAPI (1:1000, Invitrogen).

### Imaging

Sections were imaged with a wide-field scanning microscope (Olympus VS120) at 20x magnification for immunofluorescence and at 40x magnification for trichrome stained slides. Imaging large fields of view was achieved by imaging multiple, smaller images and combing them to a larger overview, which were automatically aligned by the imaging software (Olympus). All imaging was done at the Harvard Medical School Neurobiology Imaging Facility at the Department of Neurobiology (NIH NINDS P30 Core Center Grant NS072030).

### Image analysis

Original image formats were imported into FIJI using Bio-formats importer. Only the parasagittal sections spanning the distance between the olfactory turbinates and nasion were selected for analysis. The 2D, 10um sagittal images restricts data interpretations to the anterior-posterior and dorsal-ventral axis of the snout. Serial sections provide an outline to the medial-lateral aspects of the snout. Descriptions of cellular differentiation in the midface were determined by comparing anatomical location, cell morphology, cell histology, and molecular markers in serial sections within the same specimen. Levels of molecular marker Pax7 and Sox9 were quantified in the midface and then calculated as percent of DAPI-positive cells, as a measure for undifferentiated ectomesenchyme and differentiated precursor cells, respectively. Proliferating cells (PH3-positive) within each tissue section were quantified in the midface region as the percent of DAPI-positive cells. Similarly, gene expression of BMP (pSMAD1/5/9) within the midface region was quantified in each parasagittal section. Specimen data at each stage were an average of three serial sections at the parasagittal plane. Species data at each stage were an average of three specimens.

### Mouse mutant analysis

Mouse lines in this study were previously described in (Huycke et al. 2019) and briefly summarized here. Mice with a floxed *Noggin* (Stafford et al. 2011) and *Gremlin1* (Gazzerro et al. 2007) were gifted by Richard Harland (UC Berkeley), and *Dermo1-cre* (Yu et al. 2003), *Wnt1-cre* (Lewis et al. 2013) mice were obtained from JAX (stocks #008712, #022501). *Wnt1-cre* were bred to mice homozygous for both *Nog* and *Grem1* conditional alleles. The resultant mice hemizygous for all alleles were thus maintained on a mixed FVB, C57BL/6J background and crossed to homozygous floxed mice to generate embryos. Dermo1-cre mice were crossed to *Nog* and *Grem1* floxed mutants and bred in a similar manner.

### Statistical analysis

Results of midfacial length differences are expressed as mean difference between Dermo1-cre mutants and Dermo1-cre controls or Wnt1-cre mutants and Wnt1-cre controls. Two-tailed student’s t-test was used to measure statistically significant differences between groups. A value of P< 0.05 was considered statistically significant.

## Notes

### Competing Interest Statement

The authors have declared no competing interest.

