## Supplementary figures and images for "BMP signaling underlies the craniofacial heterochrony in phyllostomid bats, a hyperdiverse mammal group"

### Supplemental Fig 1

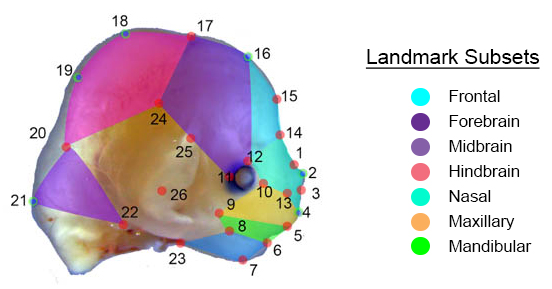

### Supplemental Fig 2

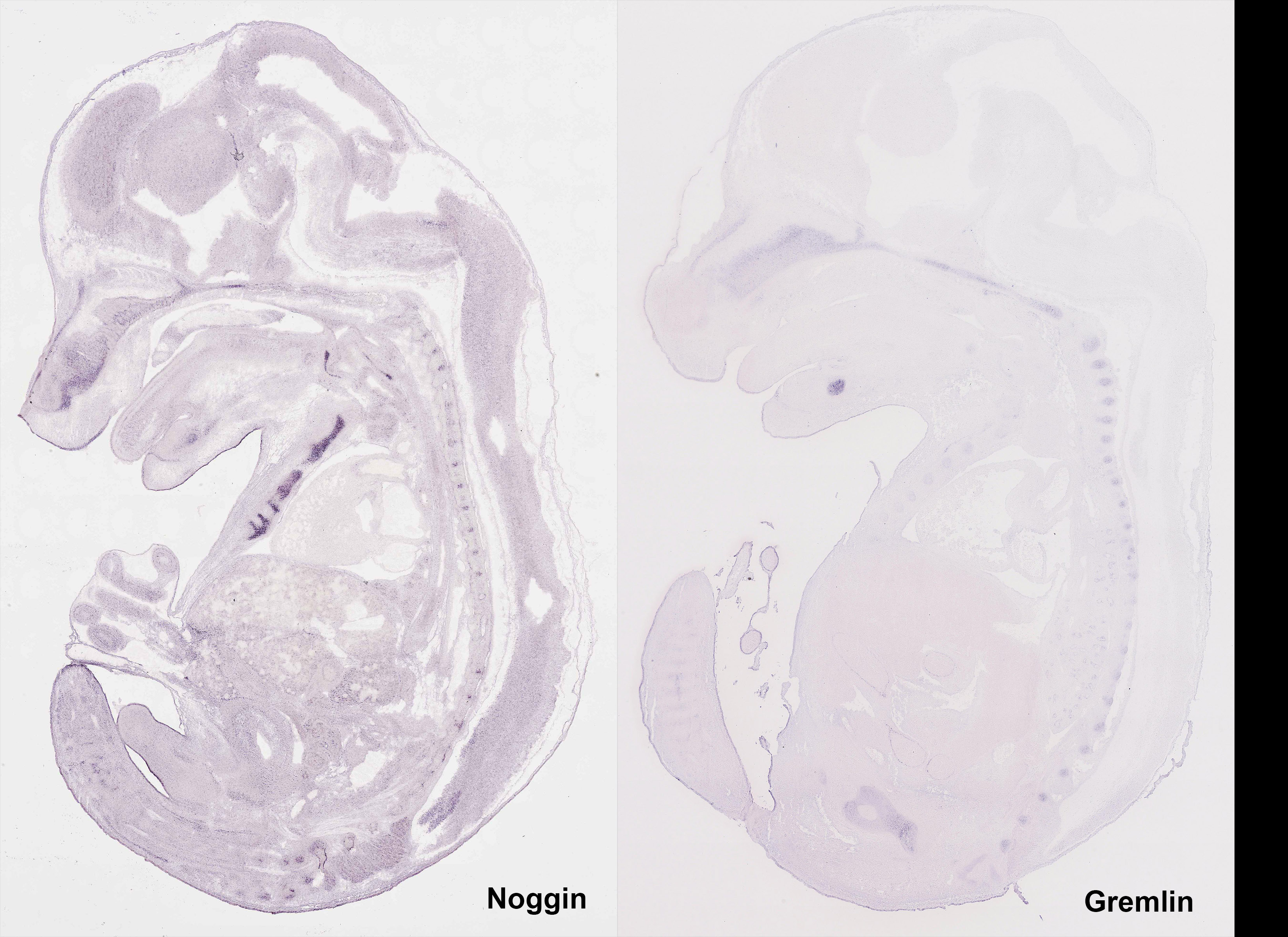

### Supplemental Fig 3

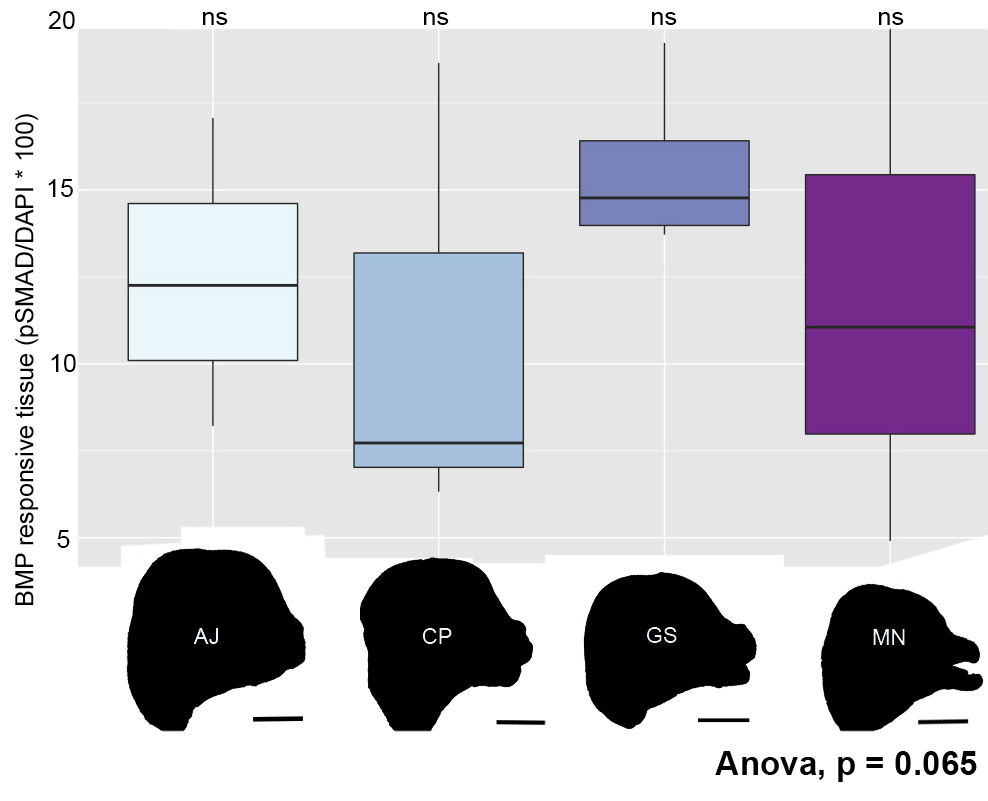
