## Supplemental figure legend for "BMP signaling underlies the craniofacial heterochrony in phyllostomid bats, a hyperdiverse mammal group"

### SUPPLEMENTAL FIGURE LEGENDS

#### S1. 2D Landmarks.

Lateral view of surface landmarks with anatomical regions identified by landmark subsets.

Table Biological definitions of all landmarks and landmark subset categories. Biological definitions for all landmarks found in Figure S1.

| LM # | Biological definition | Landmark subset |
| --- | --- | --- |
| 1 | anterior nasal | nasal |
| 2 | tip of the nose | nasal |
| 3 | nasal aperture | nasal |
| 4 | dorsal border of the maxilla and nasal aperture | nasal, maxillary |
| 5 | rostral most end of the maxilla | maxillary, dentary |
| 6 | ventral aspect of the maxilla | dentary, mandibular |
| 7 | ventral aspect of the mandible | mandibular |
| 8 | corner of developing mouth | dentary, mandibular |
| 9 | posterior border of the maxilla | maxillary, dentary |
| 10 | caudal end of nasolacrimal duct | nasal, maxillary, eye |
| 11 | posterior border of the eye | forebrain, eye |
| 12 | dorso-caudal point of lateral nasal | nasal, frontal, forebrain, eye |
| 13 | dorsal border between maxilla and lateral nasal | nasal, maxillary |
| 14 | olfactory lobe | nasal, frontal |
| 15 | rostral most point of the frontal | frontal |
| 16 | dorsal most point of the frontal | frontal, forebrain |
| 17 | border between the forebrain and midbrain | forebrain, midbrain |
| 18 | dorsal edge of the midbrain | midbrain |
| 19 | caudal most point of the brain | midbrain |
| 20 | fourth ventricle | midbrain, hindbrain |
| 21 | caudal most point of the head | hindbrain |
| 22 | carotid artery | hindbrain |
| 23 | angle of the mandible | mandibular |
| 24 | ventral caudal point of the forebrain | forebrain, midbrain |
| 25 | dorsal border of the ear | forebrain, ear |
| 26 | ear canal | ear |

#### S2. Noggin and Gremlin expressed during craniofacial development

Gene expression of Noggin and Gremlin. The Bmp antagonists Noggin (left) and Gremlin (right) are expressed during craniofacial development in mammals. They are localized within the midline cartilaginous nasal septum and cranial base in E14.5 mice. RNA *in situ* images were acquired from the Gene Expression Database (GXD) from Mouse Genome Informatics (informatics.jax.org). Noggin data were from MGI:4534065 and Gremlin data were from MGI:4519459 (Diez-Roux et al. 2011).

S3. Quantification of BMP at CS17
